# Screen Reveals Novel Roles for Tau, and Morphogen Gradients in Resolving an Epithelial Identity Crisis

**DOI:** 10.64898/2026.08.05.743011

**Authors:** Menna El Gammal, Olga Klipa, Fisun Hamaratoglu

**Affiliations:** Cardiff University

## Abstract

Epithelia, surrounding our organs and bodies, form the first line of defence against environmental insults such as wounding and infection. A constant cell turnover is essential for the renewal and integrity of the epithelium. The high turnover leads to high frequency of mistakes, and makes epithelial cancers, termed carcinomas, the most prominent cancer type. Hence, epithelial turnover and homeostasis need to be strictly controlled. The developing *Drosophila* wing consists of a two-layered epithelium and constitutes one of the best model organs to study epithelial development and homeostasis. Here, we describe our three-tiered screening efforts that led to the identification of new players in epithelial maintenance.

Selector genes act as location identity cards during development, and cells expressing the wrong selector genes for a specific location are eliminated from the epithelium. We experimentally modulate levels of Apterous, the dorsal identity gene, in cell clones, to generate cells destined to be eliminated. We isolated such cells along with their immediate neighbours with laser microdissection and determined differentially expressed genes compared to control patches. We screened these candidate genes for their ability to influence elimination of aberrant cells using RNA interference, identifying five proteins with novel roles in epithelial homeostasis: Tau, a neuronal protein, Shifted, a modulator of morphogen gradients, Wnt oncogene analog 4, p-element induced wimpy testis, which silences transposons and CG5567, a phosphatase predicted to be involved in glycerol biosynthesis. The next step will be to delineate their roles in the recognition and elimination of harmful cells.

## Introduction

Epithelia form protective barriers and allow the formation of functionally and structurally distinct tissues and organs. A constant cell turnover is essential for the renewal and integrity of the epithelium, as well as for responding to environmental insults, such as wounding (Baker, 2020; Fournon-Berodia I., 2025; Gudipaty and Rosenblatt, 2017; Levayer and Moreno, 2013). Recent work has revealed molecular players of epithelial remodelling, such as ERK signalling, that are conserved from flies to sea anemones to vertebrates, suggesting origins very early in the evolution of animals, potentially co- incident with the evolution of epithelia themselves (Fournon-Berodia I., 2025; Moreno et al., 2019). An imbalance in cell turnover, over-proliferation or reduced cell loss, can cause carcinomas (Gudipaty and Rosenblatt, 2017; Levayer and Moreno, 2013). Maintaining epithelial homeostasis is therefore of paramount importance, yet it remains insufficiently understood.

There are at least three similar, but distinct, processes that ensure epithelial homeostasis. *Cell competition*, first characterised in Drosophila in the 1970s, refers to the removal of “weaker” cells by their healthier neighbours (Morata, 2021). Today, we know that cells compete for space, nutrition and morphogens, and the “losers” can be pushed out of epithelium following apoptosis, or sometimes whilst still alive (Eisenhoffer et al., 2012; Levayer and Moreno, 2013; Morata, 2021). Loser mutations hinder house- keeping functions, such as ribosome biogenesis, proteostasis, or genome maintenance (Baker et al., 2019; Baumgartner et al., 2021; Langton et al., 2021; Ochi et al., 2021). A more recent term, *Epithelial Defence Against Cancer (EDAC),* refers to the intrinsic ability of normal epithelial cells to recognize and remove their transformed neighbours. Ras, Src or ErbB2-transformed cells are recognized by their healthy neighbours and pushed out of the epithelium, typically via apical extrusion. Importantly, when transformed cells or cells with loser mutations are on their own, and not in contact with normal cells, neither apical extrusion nor cell death is observed. Thus, this points to a non-cell-autonomous phenomenon where the transformed/loser cells are considered losers only in the presence of normal neighbouring cells (Tanimura and Fujita, 2020).

Finally, *interface surveillance* ensures recognition and elimination of mis-specified cells (cells whose identity and location are incoherent) from epithelia (Prasad et al., 2022). The Classen Lab demonstrated that two adjacent cells compare their fate and differentiation status (Prasad et al., 2022). Cells respond to pronounced fate differences between them by recruiting actomyosin to their shared contact surfaces, driving cell segregation between two clonal cell populations (Bielmeier et al., 2016). This is accompanied by elevated non-cell-autonomous apoptosis: the presence of apoptotic cells amongst both aberrant and normal neighboring cells. This response to relative differences in cell fate and differentiation states is surprisingly consistent; actomyosin recruitment, clone smoothening, and apoptosis are induced by mosaic manipulations of all the patterning genes and pathways (Adachi-Yamada and O’Connor, 2002; Bielmeier et al., 2016; Classen et al., 2009; Gibson and Perrimon, 2005; Klipa and Hamaratoglu, 2019; Liu et al., 2000; Milán et al., 2002; Organista and De Celis, 2013; Pallavi et al., 2012; Prasad et al., 2022; Prober and Edgar, 2000; Shen and Dahmann, 2005; Shen et al., 2010; Villa-Cuesta et al., 2007; Widmann and Dahmann, 2009). Whereas interface surveillance looks a lot like cell competition, there is a marked difference: interface surveillance efficiently eliminates aberrant cells in the absence of fitness differences and thus, in the absence of information about which of two neighboring cells is to survive. Instead, by comparing the program of the surrounding cells, interface surveillance uses spatial context to define a clone as ‘different’. As a consequence, it is not always aberrant cell clones that are eliminated; if the majority of cells in a region are mis-specified then the wild type cells are the ones eliminated (Bielmeier et al., 2016; Klipa and Hamaratoglu, 2021).

Whether the “aberrant” / “harmful” cells are recognized and eliminated the same way during cell competition vs EDAC vs interface surveillance remains insufficiently understood. One aspect known to be shared by EDAC (Ras-expressing cells) and interface surveillance, is bilateral JNK activation at clonal interfaces (Prasad et al., 2022). Notably, classical cell competition also activates JNK signalling, but not in a bilateral manner (Prasad et al., 2022). A full understanding of the players involved in each process will enable us to make a comprehensive comparison. Despite some new findings, there are still significant unanswered questions in our understanding of how mis-specified cells are recognized and eliminated. We do not know i) what exactly happens at the clone boundary, ii) how the wrong cell population is defined, iii) which signal(s) induce(s) the elimination program, or iiii) which downstream events cause apical shrinkage and JNK-dependent apoptosis of aberrant cells. Here, we describe our screening efforts to understand the molecular mechanisms underlying elimination of cells aberrant for their dorsal/ventral identities. First, we isolated cells to be eliminated and their immediate neighbours via laser microdissection. Then, we performed RNA-sequencing and determined the differentially expressed genes in and/or around aberrant cell clones. Next, we designed an RNAi screen to understand which of our candidate genes contribute to elimination. We focused on a list of 83 upregulated genes; we hypothesised that genes which are required for mis-specified cell elimination would result in clone rescue when downregulated within mis-specified clones. Two rounds of screening, one at the level of adult wing defects and the second scoring clone rescue in larval wing discs identified five new players: *shifted (shf), tau, Wnt oncogene analog 4 (Wnt4), p-element induced wimpy testis (piwi)* and *CG5567*.

## Results and Discussion

We generate aberrant cell clones by modulating the levels of *apterous (ap)*, the dorsal identity gene. *ap* gain-of-function cell clones have dorsal identity, and are therefore eliminated from the ventral compartment, while *ap* mutant clones have ventral identity, and are eliminated from the dorsal compartment. We defined three modes of elimination that showed spatial bias (Klipa and Hamaratoglu, 2019). If the aberrant patches touch D/V boundary, they get sorted into the appropriate compartment within the epithelium via neighbour exchanges (Blair et al., 1994; Klipa and Hamaratoglu, 2019; Milán et al., 2002). If not in contact with the D/V boundary, aberrant clones in the pouch area are more likely to undergo apoptosis, whereas basal extrusion is the main mode of elimination in the hinge area (Klipa and Hamaratoglu, 2019). We also demonstrated that the elimination of aberrant cells is independent of interfacial Myosin accumulation. We prevented Myosin accumulation by removing the Myosin Heavy chain gene from the entire tissue, from cells to be eliminated or specifically at the boundary, yet all three manipulations failed to rescue the elimination of aberrant cells (Klipa and Hamaratoglu, 2021).

### Single clone laser microdissection followed by RNA sequencing identifies differentially expressed genes at the normal-aberrant cell clone boundaries

To identify molecular changes accompanying the elimination of mis-specified cells, we used RNA-sequencing. Our workflow contained clone isolation by laser capture microdissection (LCM), library preparation, Illumina sequencing and finally, differential expression analysis (Figure 1A). We compared 3 types of clones: wild-type, *ap^DG8^* and Ap-expressing (Figure 1B). Since, we observed spatial bias in elimination mechanisms (Klipa, 2019), we prepared two separate samples for the *ap* mutant clones: isolated from the dorsal hinge (ap_Dh) and from the dorsal pouch (ap_Dp). Ap-expressing clones were isolated from the ventral pouch (Ap_Vp). Thus, we had 3 conditions of mis-specified clones. As reference points for each condition wild-type clones were isolated from the corresponding regions (wt_Dh, wt_Dp and wt_Vp). We also isolated a piece at the D/V compartment boundary (wt_DV) in order to define and exclude D/V boundary specific downstream targets. Overall, we had 7 conditions in triplicates (Figure 1B).

**Figure 1:**
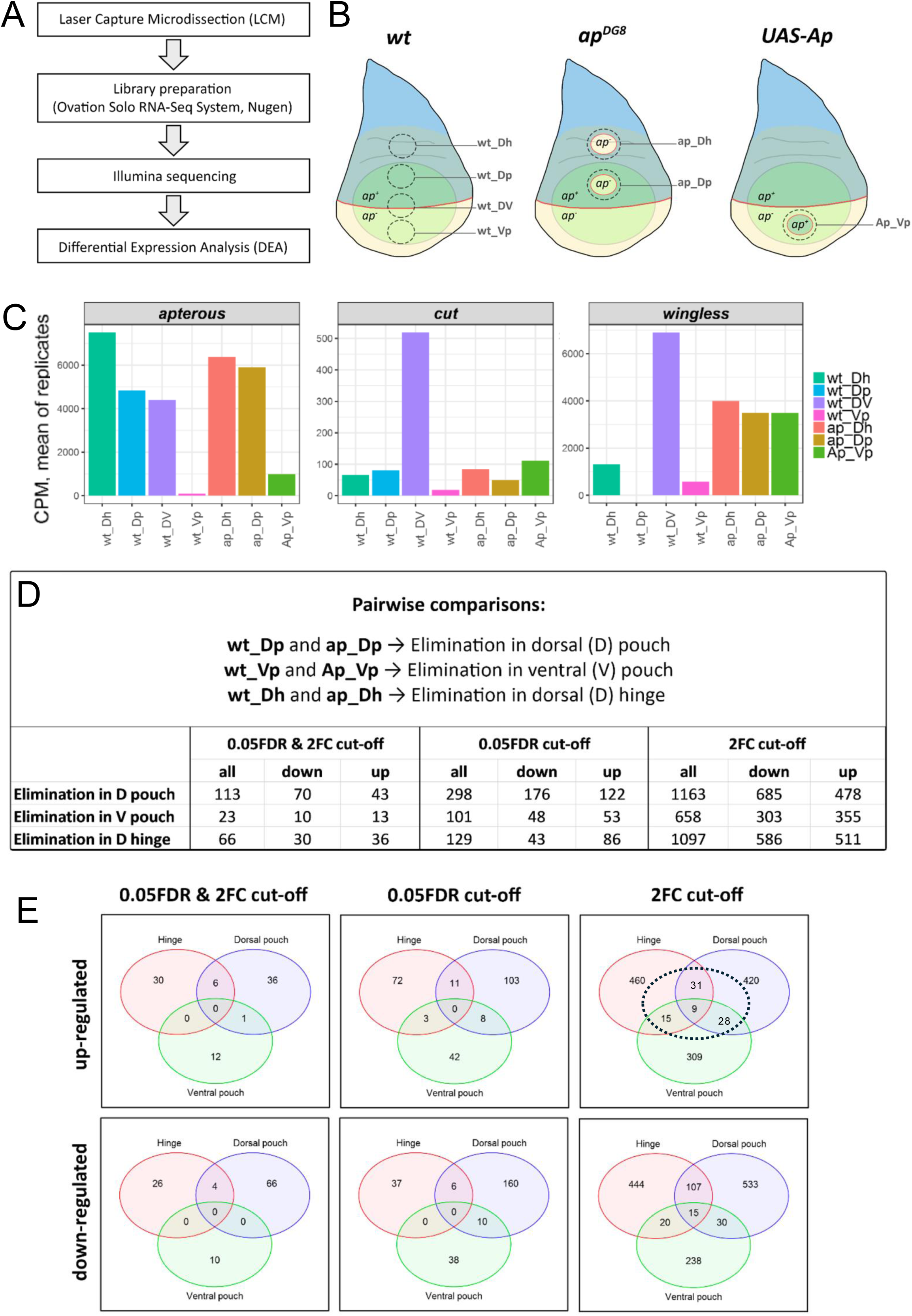
Identification of the 83 genes that were upregulated in or around aberrant cell clones. **A)** Workflow for RNA-sequencing. **B)** Schematic representation of RNA-seq samples. 7 different samples were prepared in triplicates from the indicated genotypes and 4 different regions: dorsal hinge (Dh), dorsal pouch (Dp), dorsal ventral compartment boundary (DV), and ventral pouch (Vp). Red lines represent the D/V boundary, and the dashed lines show cutting borders. For simplicity, several clones are depicted in one disc, however in most cases, one clone per disc was isolated. **C)** Expression levels of region-specific genes. Mean CPM values are shown. **D-E)** Pairwise comparisons (D), and Venn diagrams (E) showing the results of differential expression analysis of aberrant cell clones from three different regions: dorsal pouch, ventral pouch, and dorsal hinge. Results obtained with 3 different cut-off methods are displayed. Dashed circle marks the genes selected for the next screen.

We opted for LCM as the information about the clone position is preserved. Supplementary Figure 1 shows an example of the clone isolation procedure by LCM. Briefly, freshly dissected wing discs were placed on a special MMI membrane slide (Figures S1A, S1B). Clones were defined based on the GFP signal (Figures S1A’, S1B’). Note that, in order to keep clone boundary signaling, we harvested not only clonal cells but also wild-type cells in close proximity to the clone. To define the D/V compartment boundary we used vestigial QE enhancer tagged by dsRed (vg-dsRed) (Zecca and Struhl, 2007) (Figures S1A’’, S1B’’). The cutting procedure was done by MMI CellCut@ Laser Microdissection System (for further details, see Materials and Methods). The efficiency and completeness of isolation after LCM was always visually verified (Figures S1C-S1D’’). The cutting procedure resulted in local damage, hence the edges of isolated pieces were burned. To keep the clone and adjoining wildtype cells intact we defined cutting borders 2 to 3 cells away from the clone borders (Figures S1E-S1G).

In most cases 1 clone per disc was isolated. Each isolated clone consisted on average of 150 cells (min=80; max=270, cell numbers were estimated based on the measured areas and average cell density in the disc). To make libraries from single clones, we utilized Ovation Solo RNAseq System from Nugen, which is ideal for preparing libraries from 1 – 500 cells. The libraries were sequenced as single-end reads on the Illumina platform. The reads were mapped to the *Drosophila melanogaster* genome using the EMBL database. 15% – 50% of reads were uniquely aligned to exonic regions in the *Drosophila* genome. Mapped reads were principally protein coding. Gene body coverages of two replicate sets were uniform, although the samples from the third replicate set showed 3’ bias. That is most likely due to higher RNA degradation rate, as we applied more numbers of cutting rounds during LCM for the samples of this replicate. The data was normalized and batch corrected. Overall, the total number of defined genes was 16,194, among them 13,267 were protein coding. The correlations between replicates were relatively high, with Pearson’s correlation coefficients varying between R=0.857 and R=0.932.

As a quality control, we analyzed the abundance of known region-specific genes in our samples. Figure 1C shows the means of expression levels (CPM) of all replicates for *apterous, cut* and *wingless*. As expected, the dorsal selector gene *apterous* was highly expressed in the patches isolated from the dorsal compartment (Dh and Dp) and was very low in the ventral pouch. The abundance of *ap* in the mutant conditions was also relatively high, due to contribution of wild-type dorsal cells adjoining the mutant clone. The expression levels of the Notch target, *cut,* were expectedly high at the D/V boundary. Interestingly, none of conditions with mis-specified clones showed such elevated levels of *cut* expression. Expression of *wg* was similarly highest at the D/V boundary, where it is normally expressed. However, unlike *cut, wg* expression was high in our experimental conditions, reflecting its known induction at the boundary between Ap expressing and non-expressing cells (Milán et al., 2002).

Differential expression analysis (DEA) was performed using DESeq2. Pairwise comparisons of the conditions containing mis-specified cell clones with conditions containing wild-type clones from the corresponding regions revealed the up- and down- regulated genes (Figure 1D-E). We reasoned that Wg should be ectopically expressed in all scenarios, in which we had induced a boundary between cells with different *ap* expression levels. Yet, it was not detected with false discovery rate (FDR) ≤ 0.05 and fold change (FC) ≥2. Hence, we loosened our criteria for cut-off by selecting genes with either the p-adjusted value (FDR≤ 0.05), or the fold change (FC≥2). We found Wg among upregulated genes in all 3 comparisons when we used FC≥2 cut-off, but not FDR≤ 0.05, due to variability between our replicates. We decided to focus on and screen the 83 genes that were upregulated at least two-fold in at least 2 regions (dashed ellipse in Figure 1E). A list of these differentially expressed genes is presented in Table 1.

**Table 1.**

| Dorsal pouch + ventral pouch | Dorsal pouch + dorsal hinge | Ventral pouch + dorsal hinge | Ventral pouch + dorsal pouch + dorsal hinge |
| --- | --- | --- | --- |
| CG10068<br>CG1138<br>CG13894<br>CG16984<br>CG17625<br>CG3556<br>CG43277<br>CG4374<br>CG5756<br>CG9095<br>CG9657<br>CG9747<br>CheB38a<br>Cyp49a1<br>Dpck<br>E(spl)m7-HLH<br>hng3<br>l(1)G0148<br>lea<br>mamo<br>Mst87F<br>Np<br>pinta<br>piwi<br>Ret<br>roX1<br>tyn<br>Wnt4 | brk<br>CG10440<br>CG12768<br>CG13053<br>CG31807<br>CG33309<br>CG4159<br>CG43063<br>CG4573<br>CG5037<br>CG5567<br>CG5916<br>CG6023<br>CG7903<br>CG8026<br>E(spl)m2-BFM<br>Egfr<br>exu<br>fzr2<br>lama<br>magu<br>mRpL36<br>nAChRalpha1<br>otk<br>RacGAP84C<br>Ref2<br>rpr<br>sd<br>vg<br>w<br>zyd | CG10075<br>CG34001<br>CG3726<br>CG43880<br>CG44341<br>CG5210<br>CG7173<br>CG7248<br>Fbp1<br>mRpS24<br>ry<br>shf<br>SoYb<br>tau<br>TwdlF | CG12814<br>CG15756<br>CG33181<br>Cpr12A<br>rau<br>tow<br>wg<br>Wnt6<br>yellow-e3 |

### An adult wing screen eliminates half the candidates

We reasoned that if the upregulation of a gene in the aberrant cell clone has a role in clone’s subsequent elimination, reversing gene upregulation would interfere with elimination. Therefore, we carried out an RNAi screen where we downregulated our candidate genes in the mis-specified cells and scored for clone elimination efficiency. We used two independent RNAi lines per gene where possible and screened 117 *UAS-RNAi* lines. Such a screen can be performed at the larval wing disc level or by scoring adult wings because aberrant clones that were not eliminated cause developmental defects. The less effective the elimination, the more severe the defects in the wing (Klipa and Hamaratoglu, 2019).

For our screen, we generated patches that lack Apterous activity by overexpressing its negative regulator *dLMO* using the flp-out technique (Suppl. Figure 2A-C). *dLMO*- expressing clones behave in an identical manner to the *ap* loss-of-function clones (Klipa and Hamaratoglu, 2019; Milán et al., 2002). To carry out an effective genetic screen, we first established a collection of reliable and consistent controls. Aberrant cell elimination can be impaired by blocking apoptosis and JNK signalling (Milán et al., 2002; Prasad et al., 2022). We tested several transgenic fly lines, namely two lines expressing the apoptosis inhibitor p35 on different chromosomes and two constructs that can inhibit JNK signalling: a *BSK^DN^* line, and a *puckered (puc)* overexpression line (Figure 2B-C). Those were crossed to our *dLMO*-overexpressing master stock (Suppl. Figure 2A; contains a heat-shock inducible Flippase, a flp-out cassette driving Gal4 expression, *UAS-dLMO* and *UAS-GFP*) and tested under several different conditions. The variables were length of egg collection, heat shock timing and heat shock duration. The wings of the progeny were then scored for mild and severe defects and recorded. We found that the optimal conditions for our system were: a 16-hour egg collection, heat shock at 47-63 hours after egg laying (AEL) at 37°C for a period of 13 minutes (Figure 2A). We scored the progeny 2 weeks after egg collection, allowing enough time for all flies to eclose. We also counted the number of pupal cases and dead pupae at this stage. The control lines selected were *UAS-p35* (III) and *BSK^DN^* due to their higher reproducibility and significant clone rescue they displayed (Figure 2B, green and salmon bars). Upon *dLMO* overexpression in clones, 20% of all wings were deformed (Figure 2B, pink). Co-expression of *UAS-p35* (III) with *dLMO* doubled the wing defects to a frequency of 40%, and co-expression of *BSK^DN^* tripled the defects to a frequency of 67% (Figure 2B).

**Figure 2:**
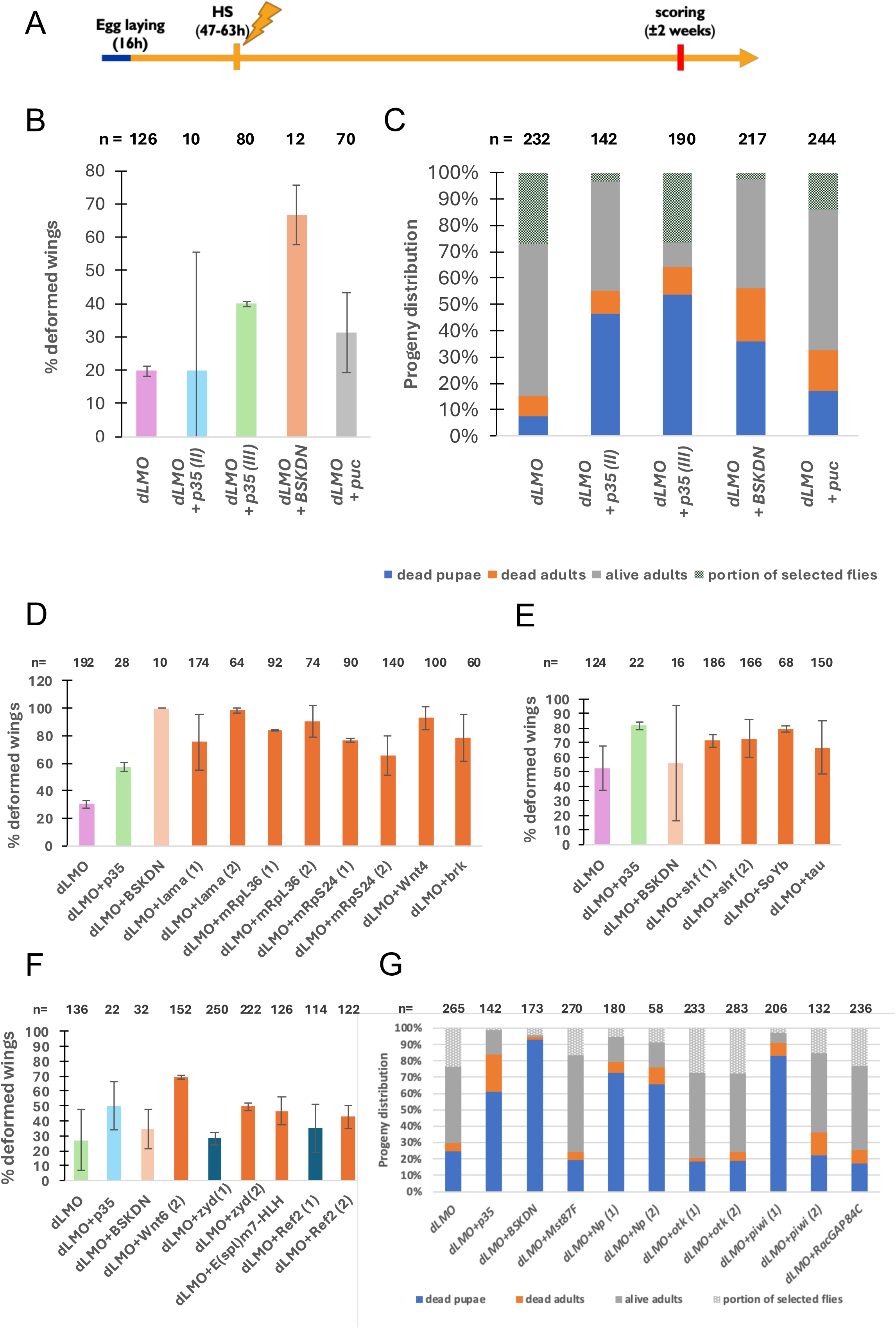
Adult wing screen identifies candidates for further analysis. **A)** Time course scheme with experimental conditions used in the adult screen. **(B)** The effect of various control genotypes on % wing deformation. n=number of wings scored of the corresponding genotypes. **(C and G)** The effects of various control (C) and experimental (G) genotypes on pupal and adult fly survival are shown. Blue blocks indicate % dead pupae; orange blocks indicate % dead adults; grey blocks indicate % non-*dLMO*-expressing alive adults; patterned blocks indicate % alive *dLMO*-expressing adults. (**D-F**) shows three experiments with hits identified shown in orange. n=number of wings scored of the corresponding genotypes. In (B-G), the averages of two replicates are shown. Error bars represent ± standard deviation.

Furthermore, blocking apoptosis or JNK signalling resulted in increased pupal and adult death (blue and orange bars in Figure 2C). About 15% of the progeny was scored dead in the presence of aberrant cell clones (*dLMO*); co-expression of *UAS-p35* or *BSK^DN^* more than tripled the frequency of progeny death. UAS-Puc had the mildest effect, but still doubled the proportion of dead pupae and adults (Figure 2C). This allows us to utilise these controls to select candidate genes based on lethality in our screen.

The screen was completed over 17 weeks. Supplementary Figures 3-5 show all the screen data by week; hits are marked in orange. To be selected as a hit, a test line should increase deformations by at least 15%, and the controls should behave as expected, that is induce significant lethality and wing deformations. It is important to compare test crosses to the control crosses from the same week as there was significant variation in the extent of deformations induced across different weeks. Crosses were performed in duplicates, and the averages are displayed. The screens where the controls didn’t behave as expected, for example screens 15 and 16, were repeated. Figures 2D- F show results of these repeat experiments. To qualify based on lethality, the test line should at least double the lethality. Based on this criterion, we shortlisted 3 lines, Np(1), Np(2) and piwi(1). All three lines induced high pupal lethality (Figure 2G, blue bars). Np(1), and Np(2) lines would not have qualified as hits based on wing deformations they induced (Suppl. Figure 5B). In total, 69 genes were tested in this assay, and we identified 44 whose upregulation in misspecified clones or their neighbours may be important for ensuring appropriate clone elimination.

### Five genes significantly rescue mis-specified clones when downregulated

After shortlisting 44 genes through the adult wing disc screen, we investigated them further at the wing disc level to verify their requirement for elimination of dLMO clones. These experiments were carried out under similar conditions as the adult screen but with a shorter egg collection period to reduce the age and size variation in wing discs (Figure 3A). Earlier clone induction typically results in the presence of larger, fewer clones, compared to later clone induction which results in smaller and more numerous clones. It is therefore important to minimize this variation for reliable interpretation and quantification of data. This screen was conducted over several different experiments, each of which is represented by a graph in Figures 3K, 4H and Supplementary Figures 6-8.

**Figure 3:**
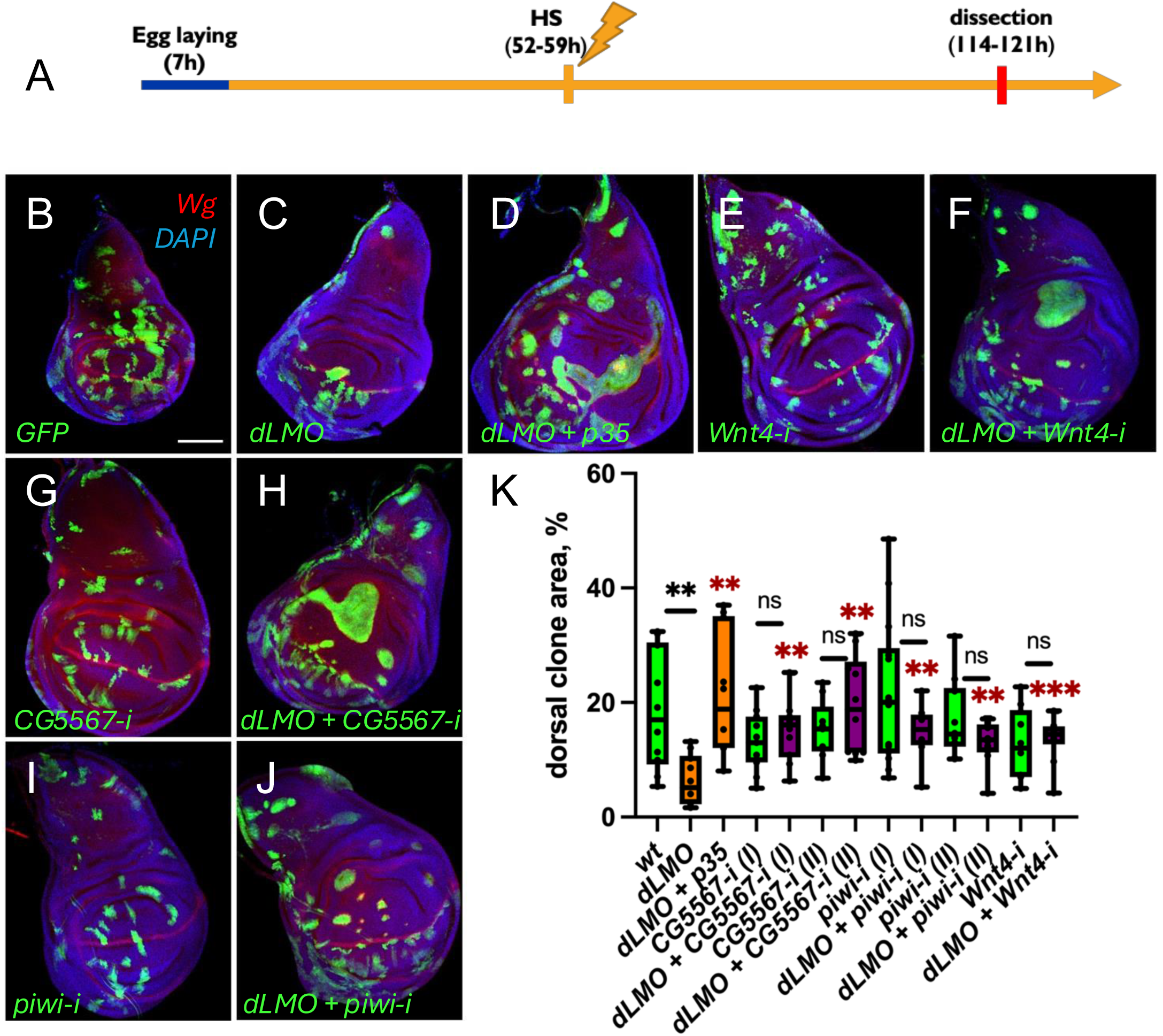
The downregulation of *CG5567*, *piwi* or *Wnt4* results in the rescue of misspecified *dLMO* clones in the dorsal compartment of the *Drosophila* wing disc. **A)** Time course scheme with experimental conditions used in the screen. **(B-J)** Representative images of wing discs expressing wild-type and *dLMO* clones both by themselves and together with *p35* (positive control) or candidate RNAi lines. Clones are marked with GFP expression (green); Wingless (Wg) antibody staining marks the D/V boundary (red); nuclei are marked with DAPI (blue). **(K)** Proportion of dorsal clone area of the various genotypes. Wild-type controls are shown in green. *dLMO* expressing lines are shown in orange. Hits are shown in purple. Number of wing discs quantified for each genotype range from 6 to 15 wing discs. The boxes represent the interquartile ranges of the data; the horizontal line within the boxes represents the median of the data; the whiskers represent the variability of the data outside the upper and lower quartiles. Significant differences were tested using a two-tailed Mann-Whitney test. Comparisons with a p-value > 0.05 were marked as “ns” (non-significant); p-value ≤ 0.05 – “ * ”; p-value ≤ 0.01 – “** ”; p-value ≤ 0.001 – “ *** ”. Scale bar in (B) represents 100µm, all images are shown at the same magnification.

**Figure 4.**
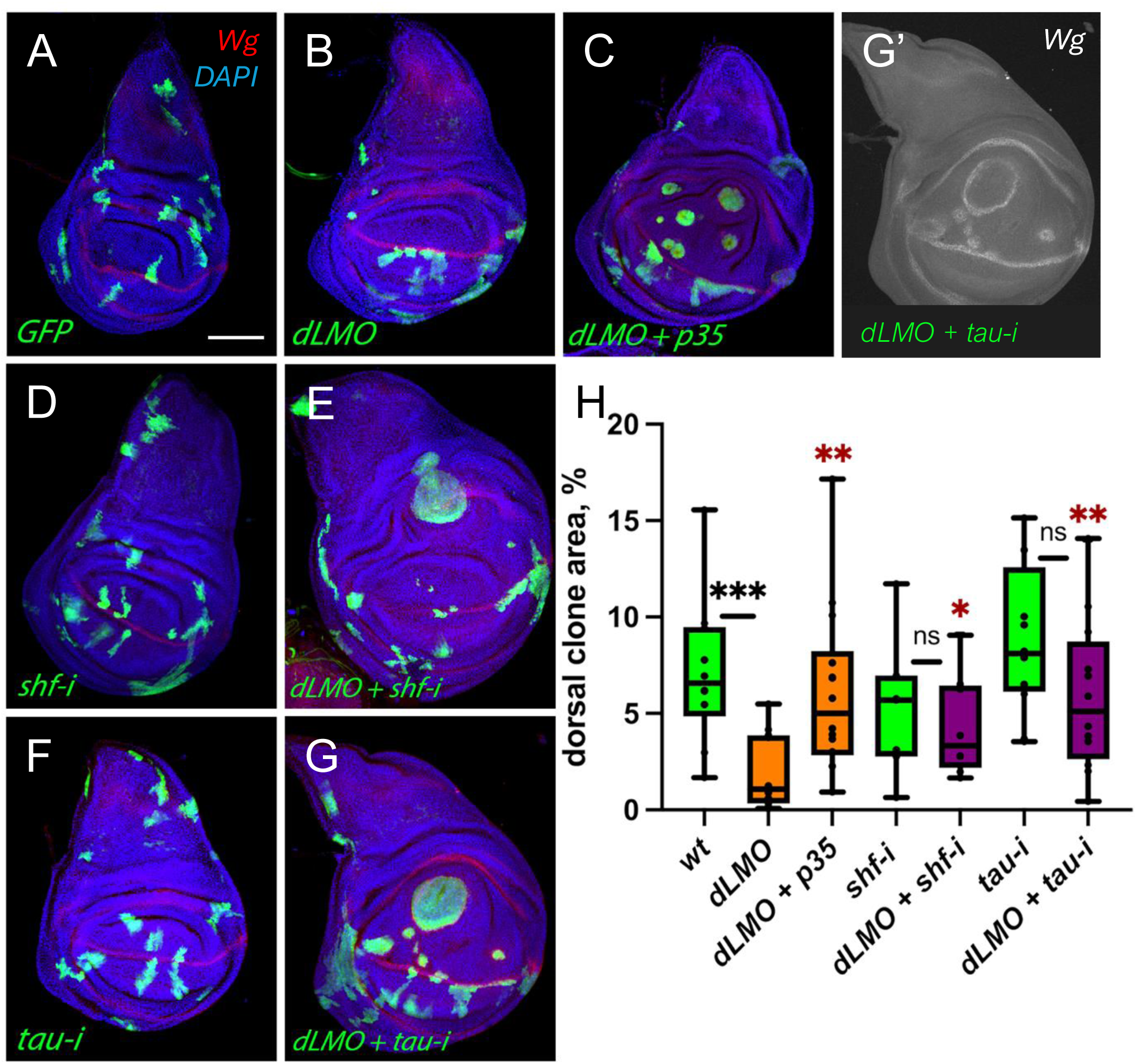
Downregulation of *tau* and *shifted* rescue the elimination of misspecified *dLMO* clones from the dorsal wing disc. A-G) Representative images of wing discs expressing wild-type and *dLMO* clones both by themselves and together with *p35* (positive control) or candidate RNAi lines. Clones are marked with GFP expression (green); Wg antibody staining marks the D/V boundary (red); nuclei are marked with DAPI (blue). **G’)** Wg channel in gray for the disc shown in G. Wg is induced around the dLMO clones in the dorsal compartment. **H)** Proportion of dorsal clone area of the various genotypes. Wild-type controls are shown in green. *dLMO* expressing lines are shown in orange. Hits are shown in purple. Number of wing discs quantified for each genotype range from 6 to 15 wing discs. The boxes represent the interquartile ranges of the data; the horizontal line within the boxes represents the median of the data; the whiskers represent the variability of the data outside the upper and lower quartiles. Significant differences were tested using a two-tailed Mann-Whitney test. Comparisons with a p-value > 0.05 were marked as “ns” (non-significant); p-value ≤ 0.05 – “ * ”; p-value ≤ 0.01 – “ ** ”; p-value ≤ 0.001 – “ *** ”. Scale bar represents 100µm, and applies to all images.

Figures 3 and 4 show representative images of wing discs for experiments involving the five hits we identified. GFP labelled control clones had irregular shapes and were found in both dorsal and ventral compartments (Figures 3B and 4A). As anticipated, *dLMO-* expressing cell clones were eliminated from dorsal compartments (Figures 3C vs 3B, 4B vs 4A), and co-expression of p35 led to significant rescue (Figures 3D, and 4C). The rescued clones were circular in shape. To quantify the rescue potential, total dorsal areas and dorsal GFP+ clone areas were measured in 6-15 wing discs for each genotype using ImageJ, from which we calculated the percentage of the dorsal compartment taken up by clones (dorsal clone area %). Black asterisks in Figures 3K and 4H indicate that *dLMO* clones occupy significantly smaller dorsal area compared to the control *GFP* clones; red asterisks display comparisons with *dLMO*.

We reasoned that for an *RNAi* line to constitute a hit, it must significantly lessen the effects of *dLMO* on dorsal clone area retention and must not by itself affect the overall dynamics of cell survival or morphology. Thus, the observed rescue of *dLMO* clones when co-expressed with *X-RNAi* must be specifically due to the gene’s role in the elimination of mis-specified cells rather than due to general cell survival or proliferation effects triggered by the *RNAi* line itself. Therefore, the percentage dorsal area occupied by *dLMO* + *candidate RNAi* clones was compared to *dLMO* alone (red asterisks) (Figures 3K, 4H, S6-8), representing the effect of the given *RNAi* line on *dLMO* clone elimination. The percentage dorsal area occupied by *dLMO* + *candidate RNAi* clones was also compared to *candidate*-*RNAi* alone (black asterisks). In cases of strong rescue, the *candidate RNAi* reverted the effect of dLMO and the dorsal clone area occupancy was not different between *dLMO* + *candidate RNAi* and *candidate*-*RNAi* alone (Figures 3K, 4H). Finally, we only selected hits from experiments in which the controls behaved as expected: *dLMO*-only discs must have a significantly lower % dorsal clone area compared to discs with GFP-expressing clones, and *dLMO + p35* discs must have a significantly higher % dorsal clone area compared to *dLMO*-only discs. Statistical differences were calculated using a two-tailed Mann-Whitney test. In this way, we were able to identify five genes whose knockdown results in a significant amount of dorsal clone rescue: *shifted, tau, Wnt4, piwi* and *CG5567*. Importantly, single *Wnt4-i* (Figure 3E), *CG5567-i* (Figure 3G), *piwi-i* (Figure 3I), *shf-i* (Figure 4D), *or tau-i* (Figure 4F), lines influence neither the morphology of the clones nor their abundance. The proportion of mis-specified clone retention in samples co-expressing *CG5567-i*, *piwi-i*, *Wnt4-i* (Figure 3)*, shf-i or tau-i* (Figure 4) along with *dLMO* is significantly higher than those expressing *dLMO* alone. For example, discs expressing *dLMO + tau-i* retained on average 4% more clonal area than *dLMO*-alone samples (Figure 4H). Strikingly, there is no significant difference in clone area in discs co-expressing *CG5567-i, piwi-i, Wnt4-i* (Figure 3)*, shf-i or tau-i* (Figure 4) along with *dLMO* (purple bars) when compared to their respective control expressing *candidate-RNAi* alone (green bars). This indicates that the *RNAi* lines effectively negated the influence of *dLMO* on clones. Whereas the rescue was impressive at the level of clone retention, the rescued clones were abnormally large and round (Figures 3F, 3H, 3J, 4E, 4G). Ectopic Wingless signal was visible around all rescued clones (Figure 3G’). The fact that the rescued clones sorted out, most likely merged with each other, and that Wg was induced at their boundaries all indicate that these cells are still recognized as “aberrant” by their normal neighbours. Thus, we conclude that all five genes must act downstream of the recognition step and upstream of the elimination step.

Our future work will focus on delineating these players in a pathway or multiple pathways of clone elimination. We will also determine whether any of these players act upstream or downstream of bilateral JNK activation.

## MATERIALS AND METHODS

### Fly Husbandry and *Drosophila* stocks

The flies were raised on cornmeal media (50L of media contains 354g agar (BPT Drewitt), 1.81kg maize flour (BPT Drewitt), 1.875kg dextrose (SLS, Cat No: FLY1156), 0,875kg dry yeast (BPT Drewitt), 111g nipagin dissolved in 1.3L EtOH, 175ml proprionic acid (Fisher, 10634622)) and crosses were kept in a 25°C incubator. The following fly stocks were used in this study: *ap^DG8^* and *FRT^f00878^* (both described in (Bieli et al. 2015), *UAS-ap* and *UAS-dLMO* (both described in Milán and Cohen 1999), *UAS-p35* (chromosome II) and *UAS-p35* (chromosome III) (both described in Hay et al. 1994), *UAS-BSK^DN^* (III) (provided by Georg Halder of KU Leuven), *UAS-puc* (described in Martín-Blanco 1998). Candidate gene RNAi lines were obtained from the Vienna *Drosophila* Resource Center (VDRC) and are presented in Table 2 along with their VDRC IDs (Dietzl et al. 2007). endnote

**Table 2:**
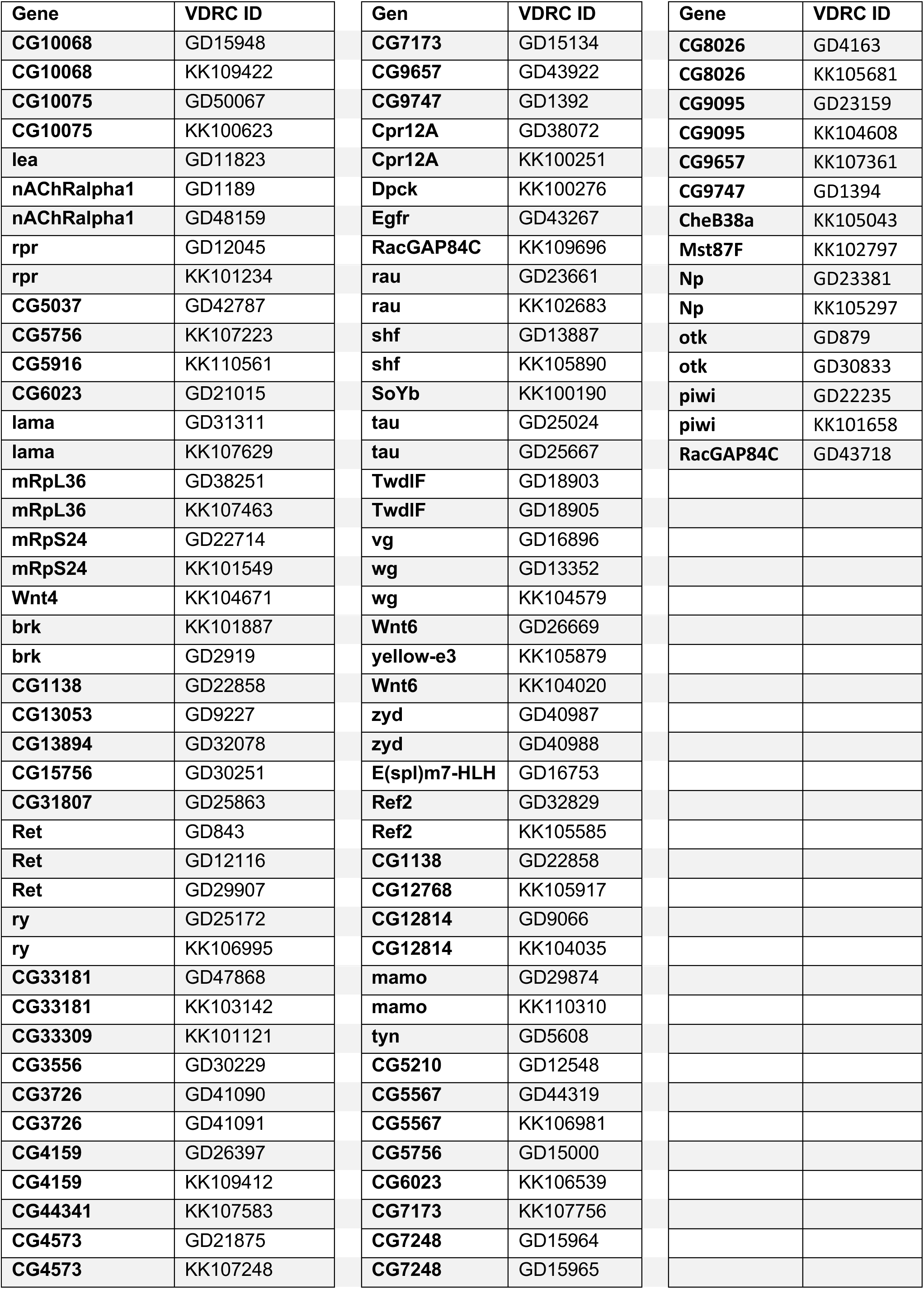
Lines used in the screen.

### Egg collection and clone induction

For adult wing screens, eggs were collected from fly crosses overnight for 16 hours at 25°C. Clone induction was temporally controlled by expressing Flippase under the control of a heat-sensitive promoter, which upon heat-shock induces excision of a stop codon surrounded by FRT sites, allowing the expression of Gal4 and UAS constructs. Heat-shock was carried out in a warm water bath 47-63 hours after egg laying (AEL). This was done at 37°C for a duration of 13 minutes. These conditions were deemed most fit after conducting a range of control optimisation experiments using various conditions and combinations. Adult wings were scored individually (for a single fly n=2) two weeks after egg collection once adults have eclosed. These experiments were carried out in duplicates. For wing disc screens, clones were induced in the same way as above, however eggs were collected from fly crosses for 7 hours only. Flippase expression was induced upon heat-shock in a warm water bath 52-59 hours AEL. This was done at 37°C for a duration of 13 minutes. Larvae were then dissected 114-121 hours after egg laying.

### Laser Microdissection and RNA-sequencing

Wing discs harbouring either *ap* loss-of-function, Ap gain-of-function or wildtype clones were dissected in cold PBS and placed on a Molecular Machines & Industries (MMI) membrane slide. The MMI Laser Microdissection system was used to excise GFP- marked clones from the tissue. Sample slides were inserted into the system along with an MMI isolation cap. Cutting was carried out at 60x magnification. The settings used were: cutting speed – 1%; focus – 50%; laser intensity – 85%. 2-6 rounds of cutting were done to achieve total isolation of the clonal region. Ovation Solo RNA-seq System kit from Nugen was used for RNA extraction and library preparation. The isolated pieces were inserted directly into the lysis buffer provided with the kit. The library preparation was carried out according to the manual. The library profiles showed good quantity and fragment size distributions, except one replicate of the wt_Vp, as revealed by Bioanalyzer. Thus, for the condition wt_Vp we had 2 replicates only. The libraries were sequenced on Illumina as single-end reads.

### Immunohistochemistry

Wing imaginal discs were prepared and stained as follows: larvae were dissected in PBS and fixed in 4% paraformaldehyde (PFA) in PBS for 30 minutes. Washes were carried out in PBS and 0.03% TritonX-100 (PBT). Samples were blocked in PBT and 2% normal donkey serum (PBTN) before they were incubated overnight in anti-Wingless (Wg) (1:2000, was deposited to the Developmental Studies Hybridoma Bank by Cohen, S.M.; DSHB Product 4D4). The secondary antibodies used were anti-mouse Alexa-Fluor 568 (Thermo Fisher). Imaginal wing discs were mounted in Vectashield mounting medium with DAPI (Vector Laboratories).

### Image acquisition and analysis

Stacked images of wing discs were acquired using a Zeiss LSM880 Airyscan confocal microscope using 20x objectives. 30-40 Z-sections were taken per image. Stacks were projected using maximum projection in ImageJ, where they were further processed and analysed. Dorsal and ventral regions were defined using Wg staining. Rescue potential (% dorsal occupation) was calculated as follows: dorsal clone area x 100/total dorsal area.

### Statistical analysis

Data processing and statistical analysis were done in GraphPad. Conditions were compared using unpaired two-samples Wilcoxon test (a.k.a. Mann-Whitney test). Comparisons with a p-value ≥ 0.05 were marked as “ns” (non-significant); p-value < 0.05 - “*”; p-value < 0.01 –“**”; p-value < 0.001 –“***”.

## Acknowledgements

We are grateful to Johann Weber, Sandra Calderon and Alexandra Dutoit of the University of Lausanne Genomic Technologies Facility for advice, data analysis and library preparation. Microscopy was performed at the Cellular Imaging Facility of the University of Lausanne and Bioimaging Research Hub of Cardiff University (RRID: SCR_022556). We thank Hoi Weeks for technical help. Thanks to Helen White-Cooper, Shijun Fang and Raham M. Rauf for comments on the manuscript.

## Figure Legends

**Figure S1.**
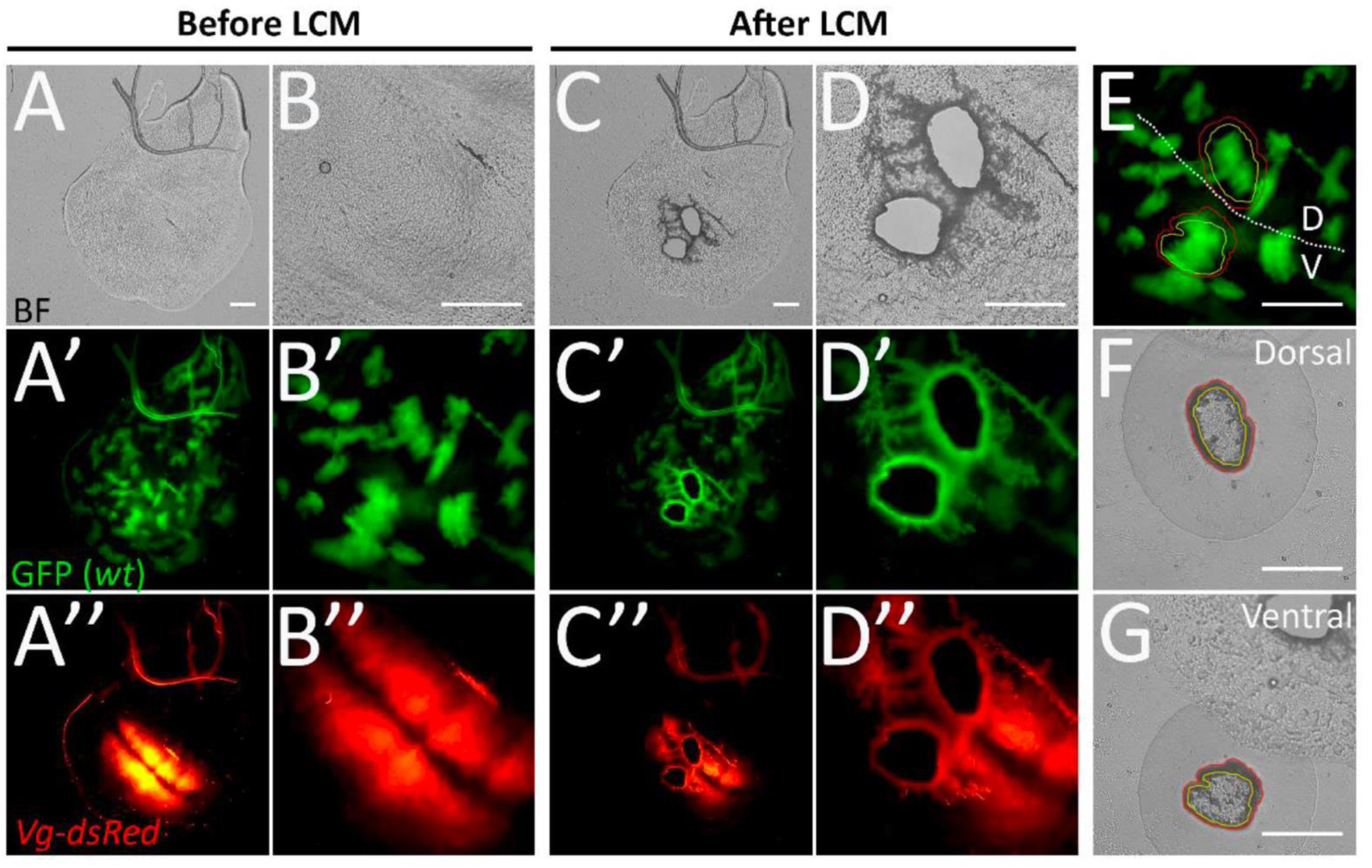
The clone isolation procedure by LCM. **(A-E)** A wing disc containing wild-type GFP-expressing clones is shown before and after LCM. **(F, G)** Isolated clones from dorsal (F) and ventral (G) compartments are shown. (A, B, C, D, F, G) – brightfield; (A’, B’, C’, D’, E) – green channel showing GFP; (A’’, B’’, C’’, D’’) – red channel showing Vg-dsRed signal. (A-B’’) show the disc before LCM at two different magnifications, low (A-A’’) and high (B-B’’). (C-D’’) show the same disc after LCM, at low (C-C’’) and high magnification (D-D’’). (E) show the same region as (B’), red lines define the contour of isolated pieces, and yellow lines demarcate the mostly intact tissue. The white dashed line represents D/V boundary. All scale bars are 50µm.

**Figure S2.**
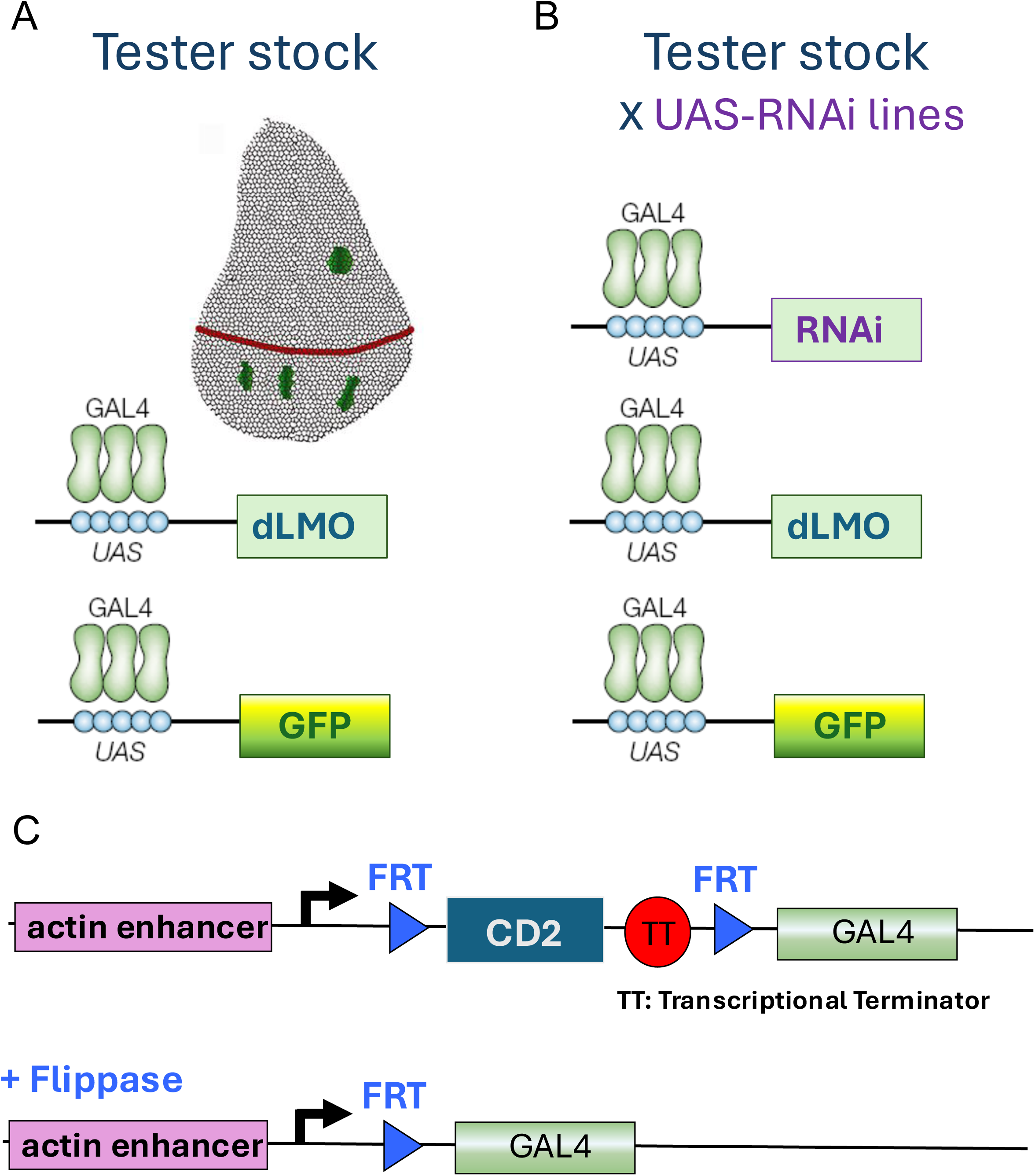
Tester stock used in the adult wing and wing disc elimination screens. **A)** A schematic of a third instar wing imaginal disc containing dLMO-expressing cell clones in green. Red line demarcates the D/V boundary. Note the circular phenotype of the remaining dLMO clone in the dorsal compartment compared to the elongated clonal shapes in the ventral compartment. Schematics below depict the binding of the GAL4 transcription factors to the UAS sites upstream of the dLMO and GFP constructs. **B)** Experimental crosses also express a UAS-candidate gene-RNAi construct as schematized. **C)** A schematic of the flp-out GAL4 construct used. Top schematic shows that there is no GAL4 expression prior to flippase expression due to the transcriptional terminator. Actin enhancer drives the expression of the marker gene CD2. Bottom: upon heat-shock driven flippase expression, in some cells FRT sites undergo recombination and the GAL4 expression is activated.

**Figure S3.**
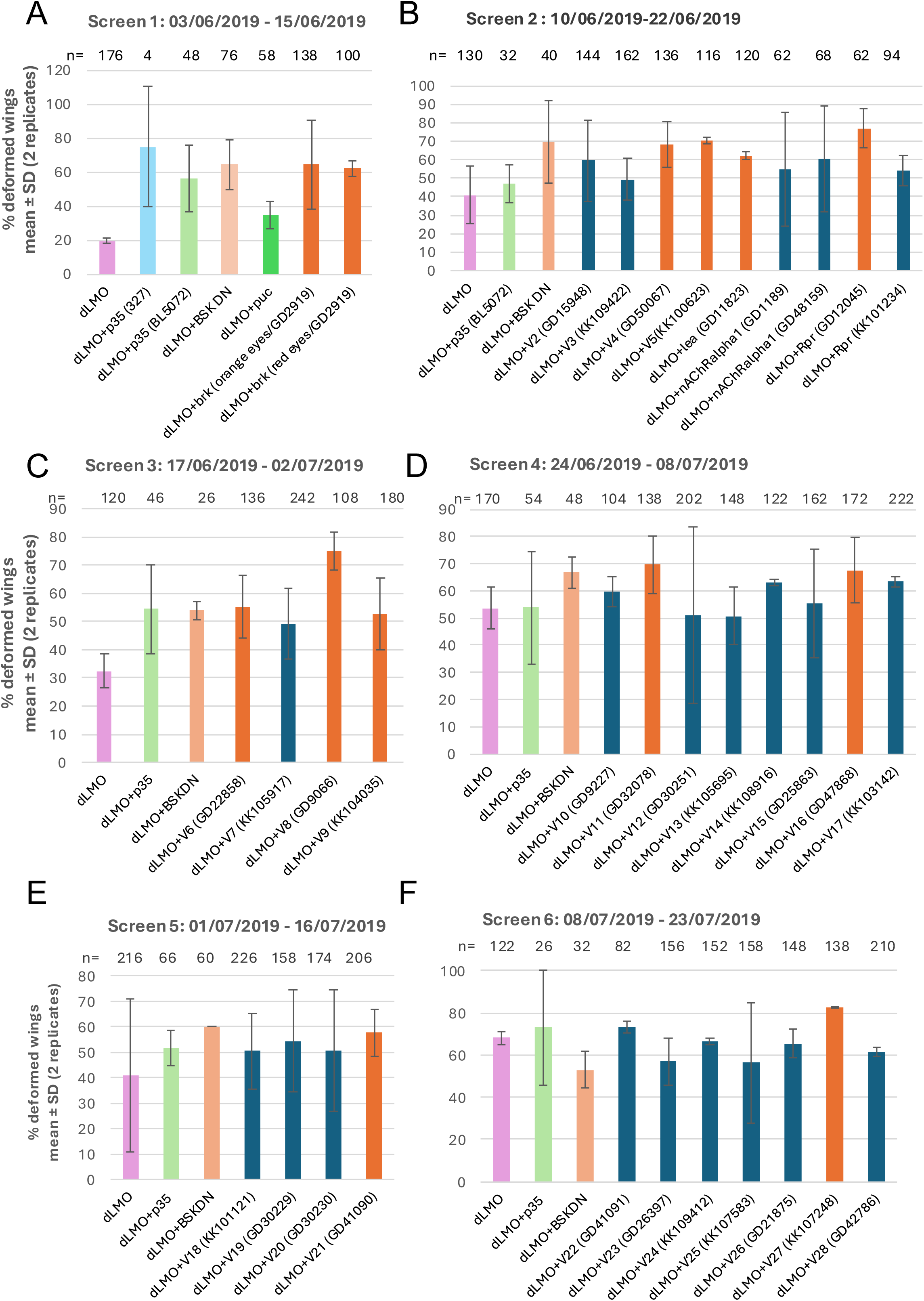
Adult wing screens. The effect of control transgenes and UAS-candidate-RNAi lines on % wing deformation. **(S3)** shows screens 1-6, **(S4)** shows screens 7-12, and **(S5)** shows screens 13-17. n indicates the number of wings scored of the corresponding genotypes. The averages of two replicates are shown. Error bars represent ± standard deviation.

**Figure S4.**
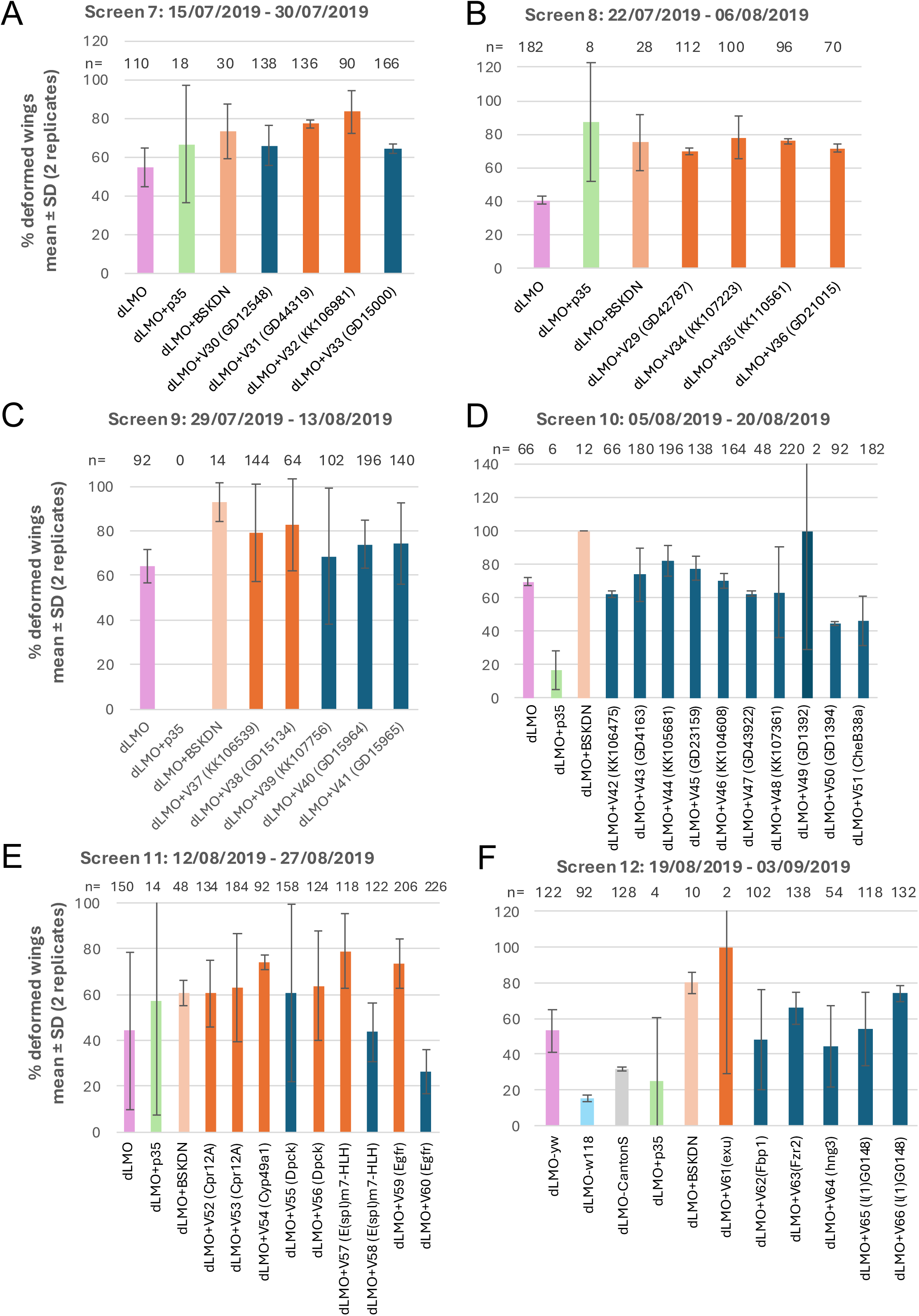
Adult wing screens. The effect of control transgenes and UAS-candidate-RNAi lines on % wing deformation. **(S3)** shows screens 1-6, **(S4)** shows screens 7-12, and **(S5)** shows screens 13-17. n indicates the number of wings scored of the corresponding genotypes. The averages of two replicates are shown. Error bars represent ± standard deviation.

**Figure S5.**
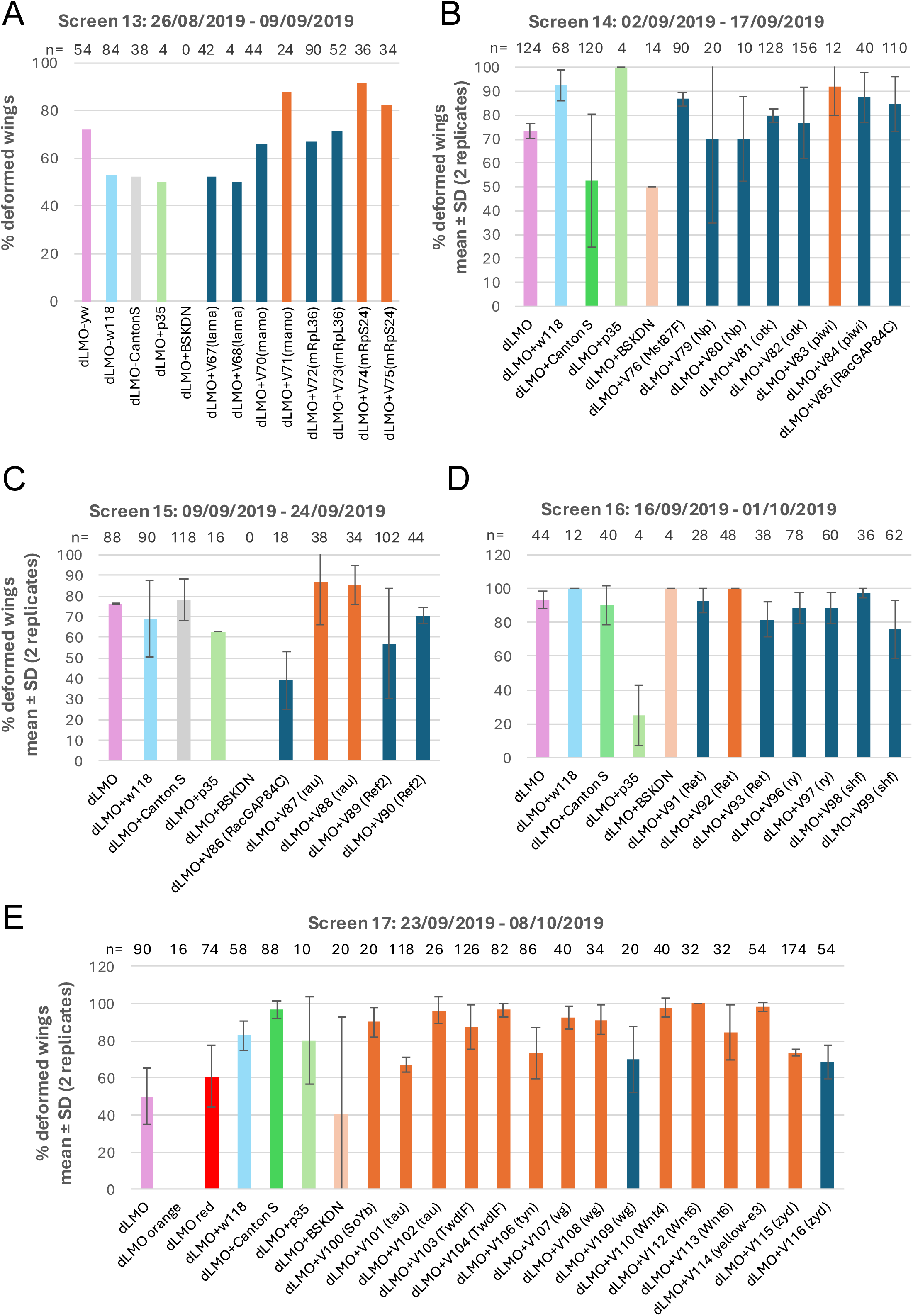
Adult wing screens. The effect of control transgenes and UAS-candidate-RNAi lines on % wing deformation. **(S3)** shows screens 1-6, **(S4)** shows screens 7-12, and **(S5)** shows screens 13-17. n indicates the number of wings scored of the corresponding genotypes. The averages of two replicates are shown. Error bars represent ± standard deviation.

**Figure S6.**
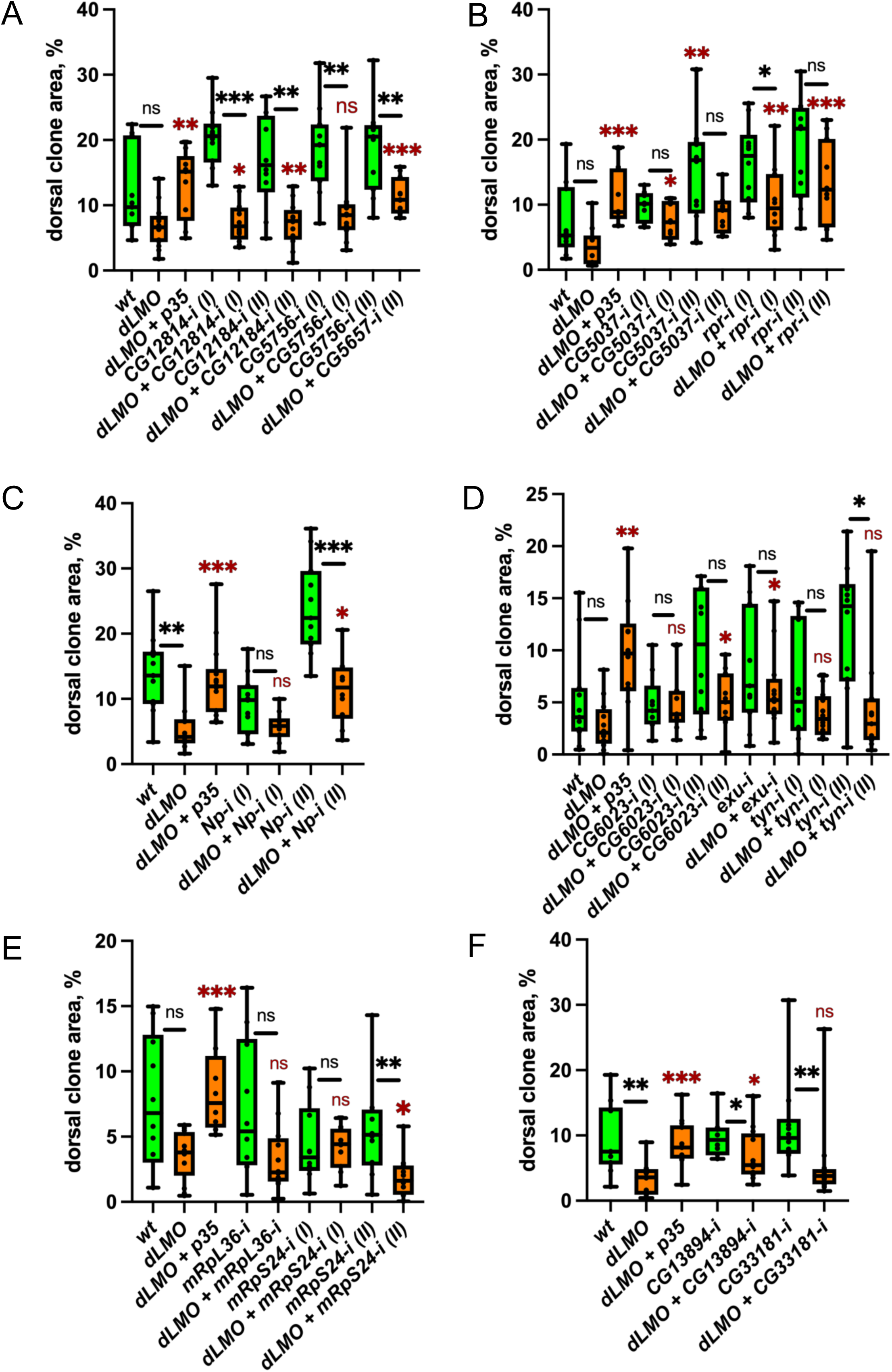
Wing disc elimination screens without hits. The graphs show proportion of dorsal clone area occupied by cell clones of the various genotypes. Wild-type controls are shown in green. *dLMO* expressing lines are shown in orange. Number of wing discs quantified for each genotype range from 6 to 15 wing discs. The boxes represent the interquartile ranges of the data; the horizontal line within the boxes represents the median of the data; the whiskers represent the variability of the data outside the upper and lower quartiles. Significant differences were tested using a two-tailed Mann-Whitney test. Comparisons with a p-value > 0.05 were marked as “ns” (non-significant); p-value ≤ 0.05 – “ * ”; p-value ≤ 0.01 – “ ** ”; p-value ≤ 0.001 – “ *** ”.

**Figure S7.**
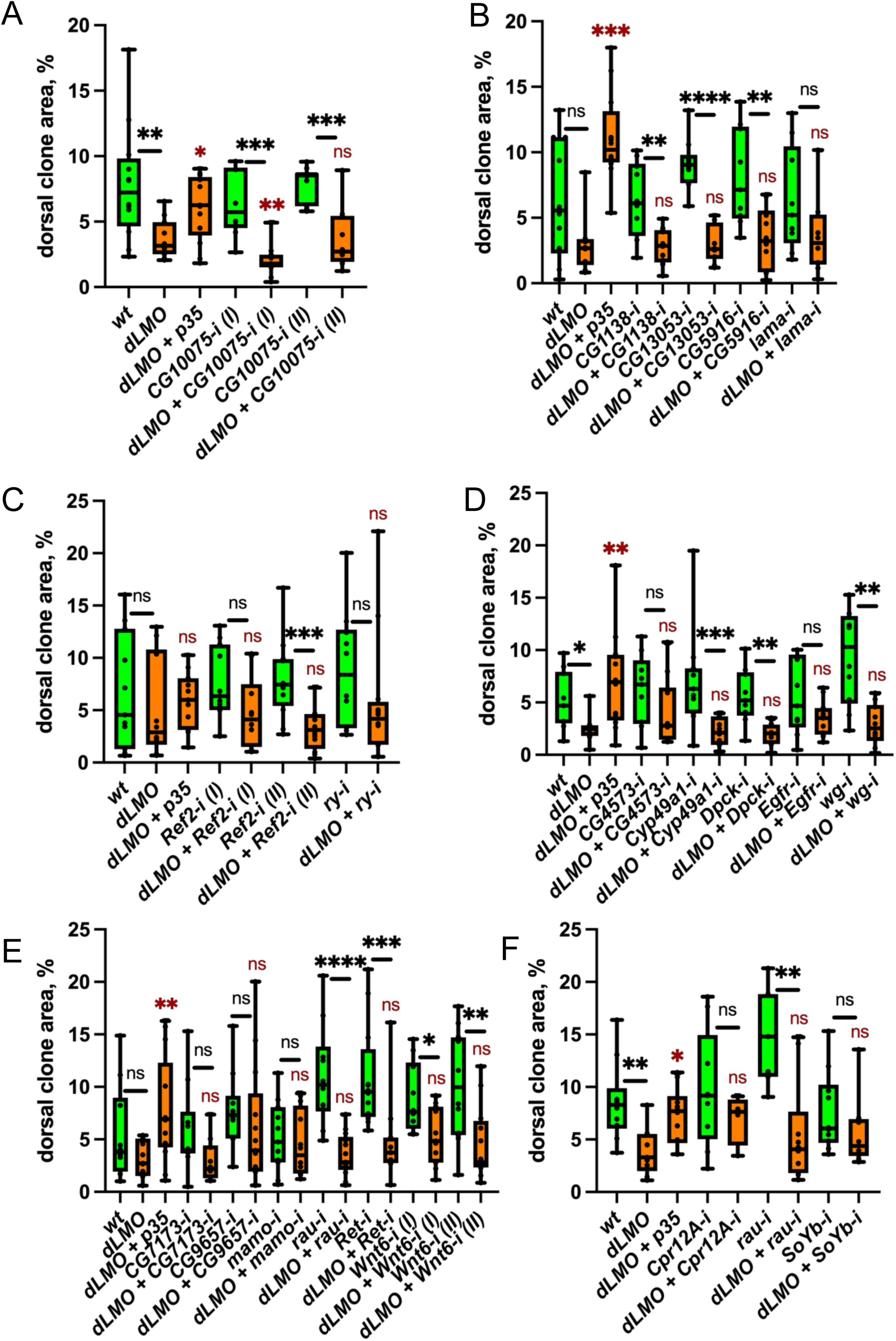
Wing disc elimination screens without hits. The graphs show proportion of dorsal clone area occupied by cell clones of the various genotypes. Wild-type controls are shown in green. *dLMO* expressing lines are shown in orange. Number of wing discs quantified for each genotype range from 6 to 15 wing discs. The boxes represent the interquartile ranges of the data; the horizontal line within the boxes represents the median of the data; the whiskers represent the variability of the data outside the upper and lower quartiles. Significant differences were tested using a two-tailed Mann-Whitney test. Comparisons with a p-value > 0.05 were marked as “ns” (non-significant); p-value ≤ 0.05 – “ * ”; p-value ≤ 0.01 – “ ** ”; p-value ≤ 0.001 – “ *** ”.

**Figure S8.**
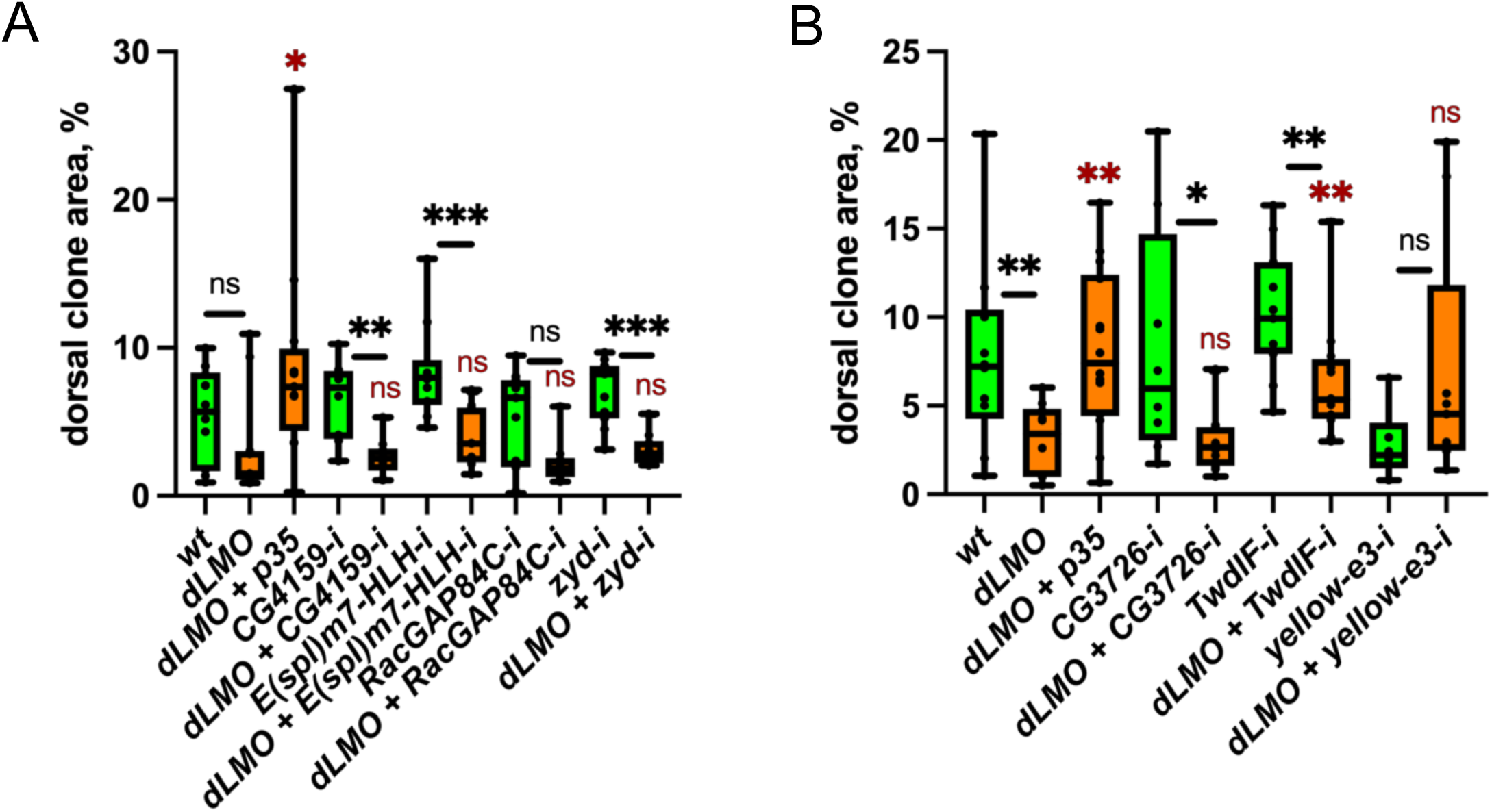
Wing disc elimination screens without hits. The graphs show proportion of dorsal clone area occupied by cell clones of the various genotypes. Wild-type controls are shown in green. *dLMO* expressing lines are shown in orange. Number of wing discs quantified for each genotype range from 6 to 15 wing discs. The boxes represent the interquartile ranges of the data; the horizontal line within the boxes represents the median of the data; the whiskers represent the variability of the data outside the upper and lower quartiles. Significant differences were tested using a two-tailed Mann-Whitney test. Comparisons with a p-value > 0.05 were marked as “ns” (non-significant); p-value ≤ 0.05 – “ * ”; p-value ≤ 0.01 – “ ** ”; p-value ≤ 0.001 – “ *** ”.

